# Improved Biosynthesis of Ethylene Glycol from Xylose in Engineered *E. coli* Utilizing Two-Stage Dynamic Control

**DOI:** 10.64898/2026.03.24.713905

**Authors:** Payel Sarkar, Shuai Li, Utsuki Yano, Jiani Chen, Michael D. Lynch

## Abstract

In this study, we employ a two-stage dynamic metabolic control strategy to enhance the NADPH dependent biosynthesis of ethylene glycol from xylose in engineered *E. coli*. We evaluated the use of metabolic valves to dynamically reduce the enzymes involved in competitive pathways which compete for substrates with ethylene glycol biosynthesis, as well as regulatory pathways aimed at increasing NADPH fluxes. The performance of our initial strains with limits in pathway expression levels was improved by the addition of competitive valves, but not by increases in NADPH flux. In contrast, improving pathway expression levels, led to strains improved significantly by our regulatory valves which improved NADPH flux, but not by the competitive valves. This is consistent with a central hypothesis that faster pathways in and of themselves can compete with other metabolic fluxes by being faster and are better aided by regulatory changes capable of change rates elsewhere in metabolism. In this case in NADPH flux. Lastly, upon scale up to fed-batch bioreactors, our optimized strain, featuring dynamic control of two regulatory valves produced 140 g/L of EG in 70 hours at 92% of the theoretical yield.

## Introduction

Ethylene glycol (EG), the simplest two-carbon polyol, is a commodity chemical utilized in polymer synthesis or directly employed as an antifreeze. ^1–4^ The global market size for EG has recenlty exceeded 30 million metric tons per year. ^5^ Currently, chemical synthesis is still the main production method. This approach relies on 3 steps: steam cracking of petrol, ethylene oxidation, and thermal hydrolysis of ethylene oxide.^6–8^ In recent years, biosynthetic approaches have been developed as alternatives, with the potential to be more environmentally friendly. ^9,10^ Current alternative routes either leverage ethanol (produced from sugars) as a feedstock with chemical conversion to EG, as well as the potential direct biosynthesis of EG from sugar feedstocks. The latter has been demonstrated in several host organisms including *E. coli* primarily producing EG from xylose.^6,11,12^

Prior studies have demonstrated the direct biosynthesis of EG using engineered *E. coli* through three main pathways. ^1,13–16^ The first leverages the Dahms pathway, which goes through the intermediate D-xylonate.^6,17–19^ The second utilizes a metabolic pathway through D-xylulose-1-phosphate, in which a ketohexokinase and aldolase are utilized to catalyze the key pathway reactions. ^19^ The third goes through D-ribulose as an intermediate and was developed based on the D-xylulose pathway. ^16^ Recently, Chae et al, reported EG biosynthesis in engineered *E. coli* leveraging the Dahms pathway relying on the overexpression of the *xylBC* genes from *C. crescentus* in addition to the *E. coli* glycolaldehyde reductase (YqhD).^6^ The fed-batch fermentation reported in this work resulted in titers of 108.2 g/L of EG with xylose as carbon source, with a yield of 0.36 g/g (0.87 mol/mol). Although this represents a reasonable titer, the high biomass levels required OD_600nm_ > 100 (estimated at > 35 gCDW/L), indicate room for further improvements with respect to the specific rate (g/gCDW*hr) of EG biosynthesis using this pathway. We have reported the use of two-stage dynamic metabolic control to improve specific production rates and titers for several products including xylitol produced from xylose. ^20–22^ The two-stage dynamic metabolic control approach, depicted in Figure 1a comprises of a growth phase, followed by a productive stationary phase, where product synthesis occurs. Phosphate depletion is used as the trigger for entry into the stationary phase as well as for the induction of genes required for product biosynthesis. Simultaneously, phosphate depletion is used to trigger proteolysis of DAS+4 degron tagged metabolic enzymes (metabolic OFF valves). ^20,21,23–25^ We sought to apply this strategy in the biosynthesis of EG, in hopes of further improving specific productivity, yield and titer.

**Figure 1.**
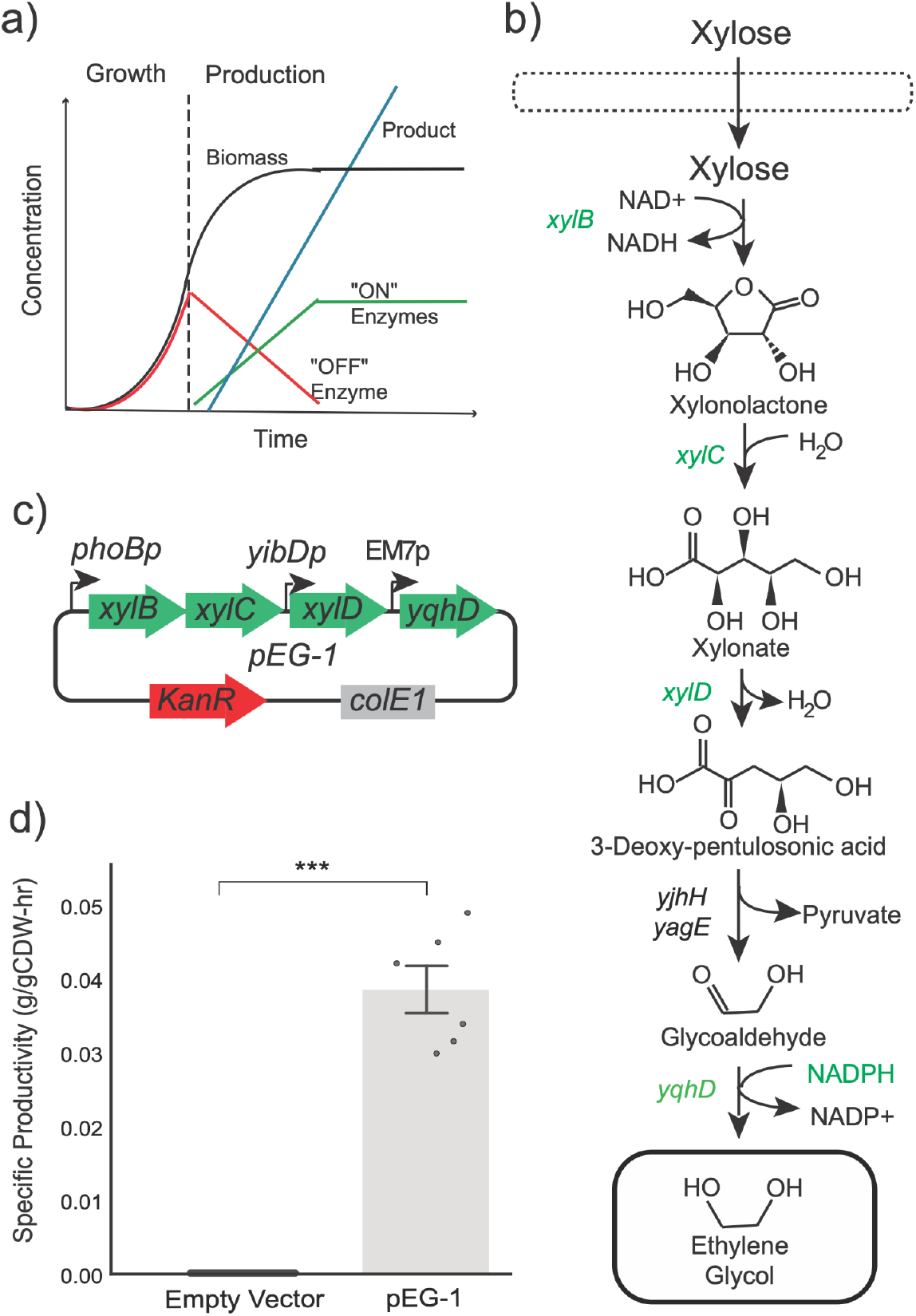
a) An overview of two stage dynamic metabolic control. Biomass levels (black line) accumulate during the first growth stage and consume a limiting nutrient (in this case phosphate), which when depleted leads to a stationary phase where product (blue line) is synthesized. Levels of key enzymes are dynamically increased due to phosphate limited induction (“ON Enzymes”, green line) or dynamically reduced (“OFF Enzymes”, red line) using synthetic metabolic valves, in this work based on controlled proteolysis. b) An overview of the production pathway for EG biosynthesis from xylose. The genes in green are overexpressed, while the black are native to *E. coli* and expressed under native control from the chromosome. Key biosynthetic enzymes include either a xylose dehydrogenase (XylB), xylonolactonase (XylC), xylonate dehydratase (XylD), endogenous enzymes 2-keto-3-deoxy D-xylonate aldolase (YjhH) or 2-keto-3-deoxy D-xylonate aldolase (YagE), and a glycoaldehyde reductase (encoded by *yqhD*). While *yqhD* native to *E. coli* it is overexpressed. c) Map of plasmid pEG-1 which enables EG biosynthesis. This plasmid bears a high copy ColE1 origin of replication and a kanamycin resistance cassette. pEG-1 contains a 2 gene operon consisting of the *xylB* and *xylC* genes driven by phosphate depletion by the phoB promoter, followed by the low phosphate inducible yibDP gene promoter driving expression of *xylD* and lastly constitutive expression of the yqhD gene via the EM7 promoter. ^50^ d) Initial specific ethylene glycol production (g/gCDW-hr) in microfermentations of the control host DLF_Z0025 with plasmid pEG-1 compared to an empty vector control (pSMART-HCKan-ev). A triple asterisk indicates a p-value <0.001, whereas a double asterisk indicates a p-value <0.01, both using a Welch’s t-test.

The application of dynamic metabolic control has proven effective in enhancing bioproduction by precisely regulating metabolic fluxes in the stationary phase. Our prior work in this area has focused on production pathways with single enzymatic steps. In these initial cases, it was not a goal to assess the interplay of pathway flux with an “optimal” metabolic network. Specifically, in this work we identify that a faster pathway in and of itself can compete with other metabolic fluxes, reducing the importance of controlling competitive fluxes and increasing the importance of metabolic changes which improve the rates of precursor synthesis. The multi-enzymatic pathway for ethylene glycol biosynthesis provides a unique opportunity to measure this behavior. By dynamically controlling both stoichiometric valves that reduce competitive fluxes and regulatory valves that increase a key cofactor flux (NADPH) in strains with low and high pathway expression levels, we observe the optimal network shift from stoichiometric to a greater dependence on regulatory changes that supply sufficient cofactor to meet the higher flux demand.

## Results

We first designed initial *E. coli* strains for two-stage EG production, using a *Dahms* pathway reliant on NADPH as the reductant as illustrated in Figure 1b. ^6,26–28^ Briefly, xylose is first oxidized to xylonate by xylose dehydrogenase (XylB) and xylonolactonase (*XylC*). Next, the conversion of xylonate to 3-Deoxy-pentulosonic acid is carried out by xylonate dehydratase (XylD). Then, native endogenous enzymes encoding 2-keto-3-deoxy D-xylonate aldolase (YjhH) or 2-keto-3-deoxy D-xylonate aldolase (YagE) convert the 3-Deoxy-pentulosonic acid to glycolaldehyde. Lastly, the NADPH-dependent aldehyde reductase (YqhD from *E. coli*) is overexpressed to reduce the glycolaldehyde to EG. ^6,14^ First, we constructed plasmid pEG-1, illustrated in Figure 1c, expressing 4 of the pathway enzymes: xylose dehydrogenase (XylB), xylonolactonase (X*ylC*), xylonate dehydratase (XylD), and glycoaldehyde reductase (YqhD). Refer to Table 1 for a list of strains and plasmids used in this study. This plasmid was transformed into strain DLF_Z0025, which is a control strain containing phosphate induction machinery but lacking any valves (degron tagged enzymes). This strain, as well as an empty vector control, was then evaluated for EG production in two stage microfermentations completed in 96 well microtiter plates. ^29^ As Figure 1d shows, overexpression of the 4 pathway enzymes indeed led to EG production (which was absent in the empty vector control) with a specific production rate of 0.034 g/gCDW-hr.

**Table 1:** Plasmids and strains used in this study.

Tables, Figures & Captions
| Table 1: Plasmids and strains used in this study |  |  |  |  |  |
| --- | --- | --- | --- | --- | --- |
| Plasmid | Insert | ori | Res | Addgene | Source |
| pSMART-HC-Kan | Empty vector | colE1 | Kan | NA | Lucigen |
| pSIM5 | Recombineering Plasmid | pSC101 | Cm | NA | <sup>46</sup> |
| pCASCADE-zwf | ugpBp-gRNA silencing zwf | p15a | Cm | 65825 | <sup>20</sup> |
| pEG-1 | phoBp-*xyIBC-yibDp-xyID-EM7p-yqhD* | colE1 | Kan | 246717 | This study |
| pEG1-nXD | phoBp-*xyIBC-EM7p-yqhD* | colE1 | Kan | 246718 | This study |
| pCOLA-xyID-1 | *yibDp-rbs1- xyID variant 1* | colA | Gent | 246719 | This study |
| pCOLA-xyID-3 | *yibDp-rbs2- xyID variant 2* | colA | Gent | 246720 | This study |
| pCOLA-xyID-4 | *yibDp-rbs1- 6xhis - xyID variant* | colA | Gent | 246721 | This study |
|  | *1-linker-HIBIT* |  |  |  |  |
| pCOLA-xyID-5 | yibDp-rbs3- 6xhis - *xyID* variant<br>*1-linker-HIBIT* | colA | Gent | 246758 | This study |

| Strain | Genotype | Source |
| --- | --- | --- |
| DLF_R004 | F-, λ-, Δ(araD-araB)567, lacZ4787(del)(::rrnB-3) , rph-1, Δ(rhaD-rhaB)568, hsdR514, ΔackA-pta, ΔpoxB, ΔpflB, ΔldhA, ΔadhE, ΔiclR, ΔarcA, ΔompT:yibDp-λR-nucA-apmR | 39 |
| DLF_Z0025 | F-, λ-, Δ(araD-araB)567, lacZ4787(del)(::rrnB-3) , rph-1, Δ(rhaD-rhaB)568, hsdR514, ΔackA-pta, ΔpoxB, ΔpflB, ΔldhA, ΔadhE, ΔiclR, ΔarcA, ΔsspB::frt, Δcas3:: ugpBp-sspB-pro-casA | 20 |
| SL_0001 | DLF_Z0025, xylA-DAS4:ampR | 20 |
| DLF_Z0763 | DLF_Z0025, udhA-DAS4:bsdR | 20 |
| PS_0001 | DLF_Z0025, udhA-DAS4:bsdR, xylA-DAS4:ampR | This study |
| PS_0002 | DLF_Z0025, fabI-DAS4:zeoR | This study |
| PS_0003 | DLF_Z0025, fabI-DAS4:zeoR, xylA-DAS4:ampR | This study |
| PS_0004 | DLF_Z0025, fabI-DAS4:zeoR, udhA-DAS4:bsdR | This study |
| PS_0005 | DLF_Z0025, fabI-DAS4:zeoR, udhA-DAS4:bsdR, xylA-DAS4:ampR | This study |
| DLF_Z1002 | DLF_Z0025, zwf-DAS4:bsdR | 20 |
| PS_0006 | DLF_Z0025, fabI-DAS4:zeoR, zwf-DAS4:bsdR | This study |
| PS_0007 | DLF_Z0025, fabI-DAS4:zeoR, zwf-DAS4:bsdR, xylA-DAS4:ampR | This study |
| PS_0008 | DLF_Z0025, fabI-DAS4:zeoR, zwf-DAS4:bsdR, udhA-DAS4:bsdR | This study |
| Res - resistance marker, ori- origin of replication,, Cm- chloramphenicol, Kan - kanamycin, Amp - ampicillin, tet - tetracycline, zeo- zeocin |  |  |

We next evaluated the impact of two key stoichiometric metabolic valves on two-stage EG production. The two valves involve the dynamic reduction of levels of the enzymes, XylA and UdhA, as illustrated in Figure 2a. XylA (xylose isomerase) leads to the catabolism of xylose through xylulose directing, competing with the production pathway for xylose. Similarly, UdhA, the soluble transhydrogenase can convert NADPH, which is required for the final step of EG biosynthesis, to NADH. Again, UdhA represents a competitive pathway for a key biosynthetic precursor. The dynamic reduction of enzymes in these competitive pathways can be expected to stoichiometrically improve EG biosynthesis, by reducing competitive substrate consumption. Importantly, we have previously reported the use of these proteolytic valves. ^20^ In the case of XylA, proteolytic degradation led to a 60% decrease in enzyme activity, and in the case of UdhA, proteolytic degradation led to a 30% decrease in enzyme activity.^20^ We have also previously reported the use of CRISPR based gene silencing in combination with proteolytic degradation to improve reductions in protein levels. In the case of both XylA and UdhA, the addition of gene silencing in the stationary phase did not improve upon reductions in activity achieved by proteolytic degradation alone. ^20,30^ Strains with either UdhA or XylA valves as well as valves in both genes were transformed with plasmid pEG-1 and assessed for EG production in two-stage microfermentations.^29^ Results are given in Figure 2b. While two-stage dynamic control over UdhA or XylA alone did not lead to improvements in EG productivity, the combination of both valves led to a > 3 fold improvement in EG specific productivity of 0.11g/gCDW-hr.

**Figure 2.**
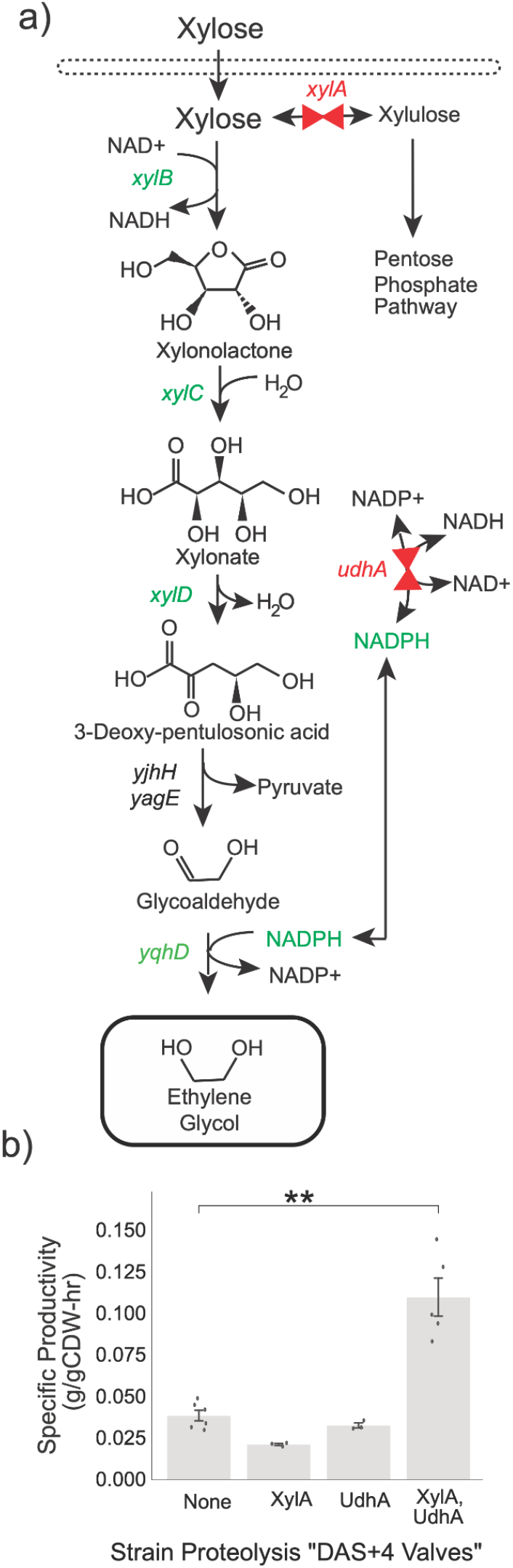
b) An overview of the production pathway for EG biosynthesis, discussed in Figure 1, in relation to two key “stoichiometric metabolic valves in the native *xylA* and *udhA* genes, shown as red valves. b) Specific EG production (g/gCDW-hr) in microfermentations of strains with plasmid pEG-1 in control host DLF_Z0025 (No valves) as well as in strains with dynamic proteolysis of XylA, UdhA or both. Note the different scale of the y-axes for panels B and C. A single asterisk indicates a p-value <0.05, whereas a double asterisk indicates a p-value <0.01, both using Welch’s t-test.

We next sought to evaluate the impact of key “regulatory valves” which we have recently reported to enhance NADPH production rates, namely valves in the key enzymes enoyl-ACP reductase (FabI) and glucose-6-phosphate dehydrogenase (Zwf). The role of these enzymes in NADPH metabolism is illustrated in Figure 3a. Briefly, acyl-ACP intermediates in fatty acid biosynthesis are known competitive inhibitors of the membrane bound transhydrogenase: PntAB. ^31–33^ Dynamic reduction of FabI activity reduces levels of acyl-ACPs, alleviating inhibition of PntAB and improving NADPH production. ^20^ We have hypothesized that Zwf has a regulatory role in controlling not only NADPH production but also NADPH/NADP+ ratios (reducing potential) and that dynamic reduction in Zwf activity leads to a decreased reducing potential and the activation of alternative NADPH synthesis pathways which operate to generate a lower NADPH/NADP+ ratio yet with higher potential NADPH fluxes. ^20^ Strains with dynamic proteolysis over FabI and Zwf were constructed along with strains with dynamic proteolysis of combinations of XylA, UdhA, FabI and Zwf. These strains were transformed with pEG-1 and assessed for EG production in two stage microfermentations. These results are given in Figure 3b. In these experiments the strain with XylA and UdhA valves still had the highest specific EG biosynthesis rate.

**Figure 3.**
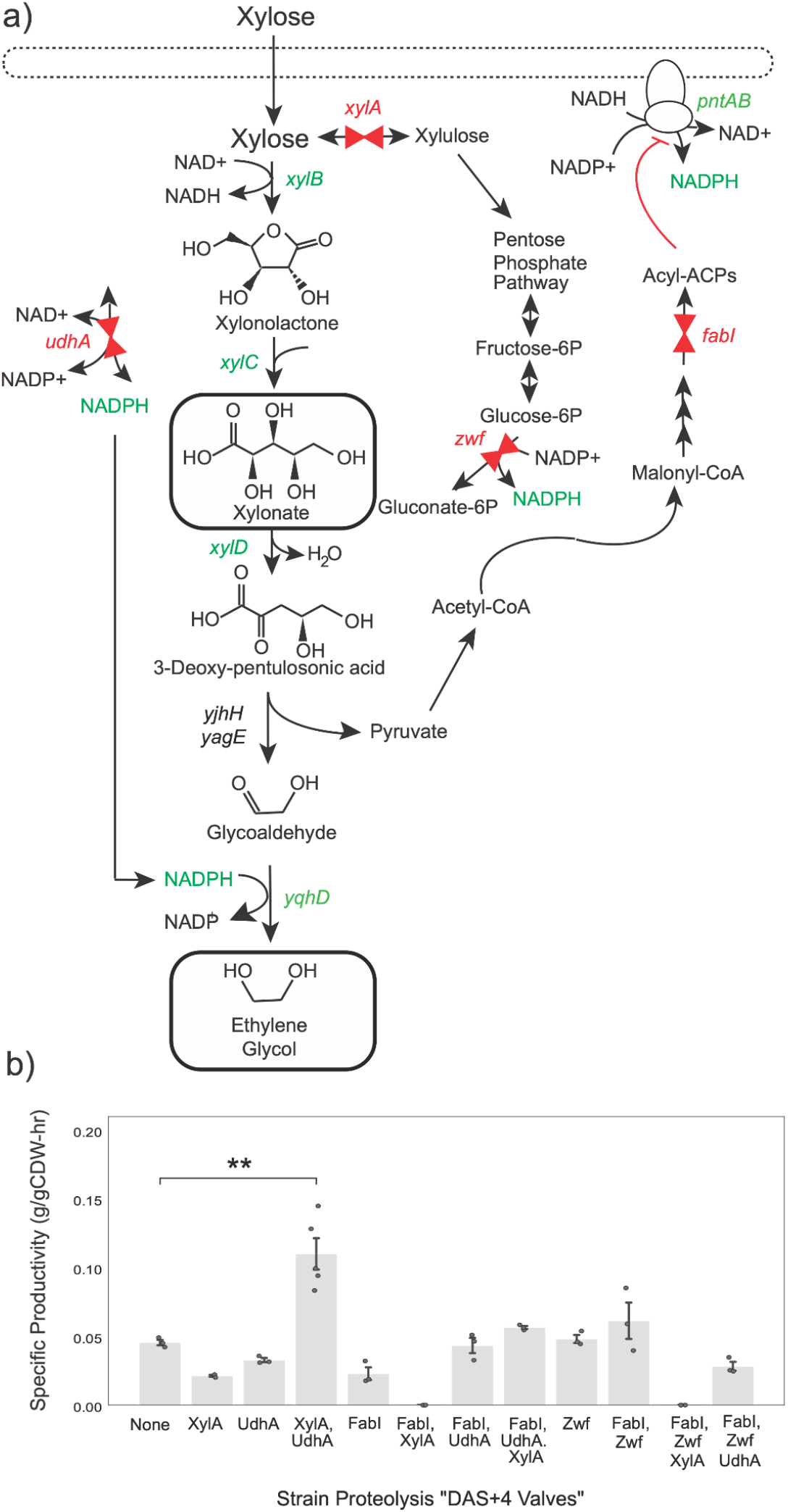
a) An overview of the production pathway for EG biosynthesis, discussed in Figure 1, in relation to the two key “stoichiometric metabolic valves in the *xylA* and *udhA* genes as well as two “regulatory” metabolic valves in the *fabI* and *zwf genes*, shown as red valves. b) Specific EG production (g/gCDW-hr) in microfermentations of strains with plasmid pEG-1 in the control host DLF_Z0025 (No valves) as well as in strains with dynamic proteolysis of XylA, UdhA, FabI and Zwf alone and or in combination. A double asterisk indicates a p-value <0.01, using a Welch’s t-test, comparing the strain with dynamic proteolysis of both FabI and UdhA to the control.

No improved strain variants were identified, and the absolute specific production rates of ∼0.1g/gCDW-hr fell lower than our expectations. The pathway enzymes chosen all have very reasonable activity (Refer to Supplemental Table S1 for reported biochemical parameters) and should be able to support higher specific production rates assuming they are reasonably overexpressed.^34–38^ To assess any potential limits in production due to low pathway enzyme expression levels, we performed SDS-PAGE analysis of plasmid pEG-1. This was performed in strain DLF_R004, which shares a similar background with DLF_Z0025, and enables two stage expression but additionally, enables autolysis and autohydrolysis of DNA and RNA simplifying expression analysis. ^39^ Results are given in Supplemental Figure S1. Briefly, three of the pathway enzymes including XylB, XylC and YqhD were obviously overexpressed in whole cell lysates as well as the soluble fraction. Densitometry estimates for the expression level (% of total protein) for each enzyme in the soluble fraction were: XylB, 8.55%, XylC 8.09%, and YqhD 33.1%. Overexpression of XylD was not observed in the soluble fraction or in whole cell lysates. We are unsure of whether XylD is not expressed at all, or just below the limit of detection for SDS-PAGE. In ether even *E. coli* does possess two native xylonate dehydratases encoded by the *yagF* and *yjhG* genes, which are capable of replacing xylD (albeit at lower activity Supplemental Table S1) and produce EG and derivatives. ^13,36,40^

Based on a lack of XylD overexpression from plasmid pEG-1, we moved to create new plasmids for improved pathway expression. These plasmids are illustrated in Figure 4a. First as it was not expressing, we removed the *xylD* open reading frame from pEG-1, to make pEG1-nXD (“no xylD”). First we confirmed that the three enzymes were still successfully overexpressed XylB, XylC and YqhD were still successfully overexpressed from pEG1-nXD, after removal of the *xylD* gene cassette. These results are shown in Supplemental Figure S2, and confirm robust overexpression in whole cell lysates as follows: XylB 10.3%, XylC 7.1%, and YqhD 19.6%. In parallel, we constructed independent plasmids for expression of several *xylD* gene variants with their own low phosphate inducible promoter in a medium copy pCOLA backbone. Variants included 2 reported isoforms, a few ribosomal binding alternatives and variants with N-terminal 6x histidine tags and C-terminal HIBIT tags (Supplemental Table S4). The latter were constructed in the case we needed to measure expression levels below the limit of detection by SDS-PAGE.^37,41^ Again, we evaluated the expression of our various *xylD* expression constructs using strain DLF_R004. These results show in Supplemental Figure S3, confirm expression in whole cell lysates. Densitometry led to rough estimates of expression level for each xylD variant, as follows: variant 1: 19.1%, variant 3: 6.82%, variant 4: 1.5% and variant 5: 7.94 %.

**Figure 4.**
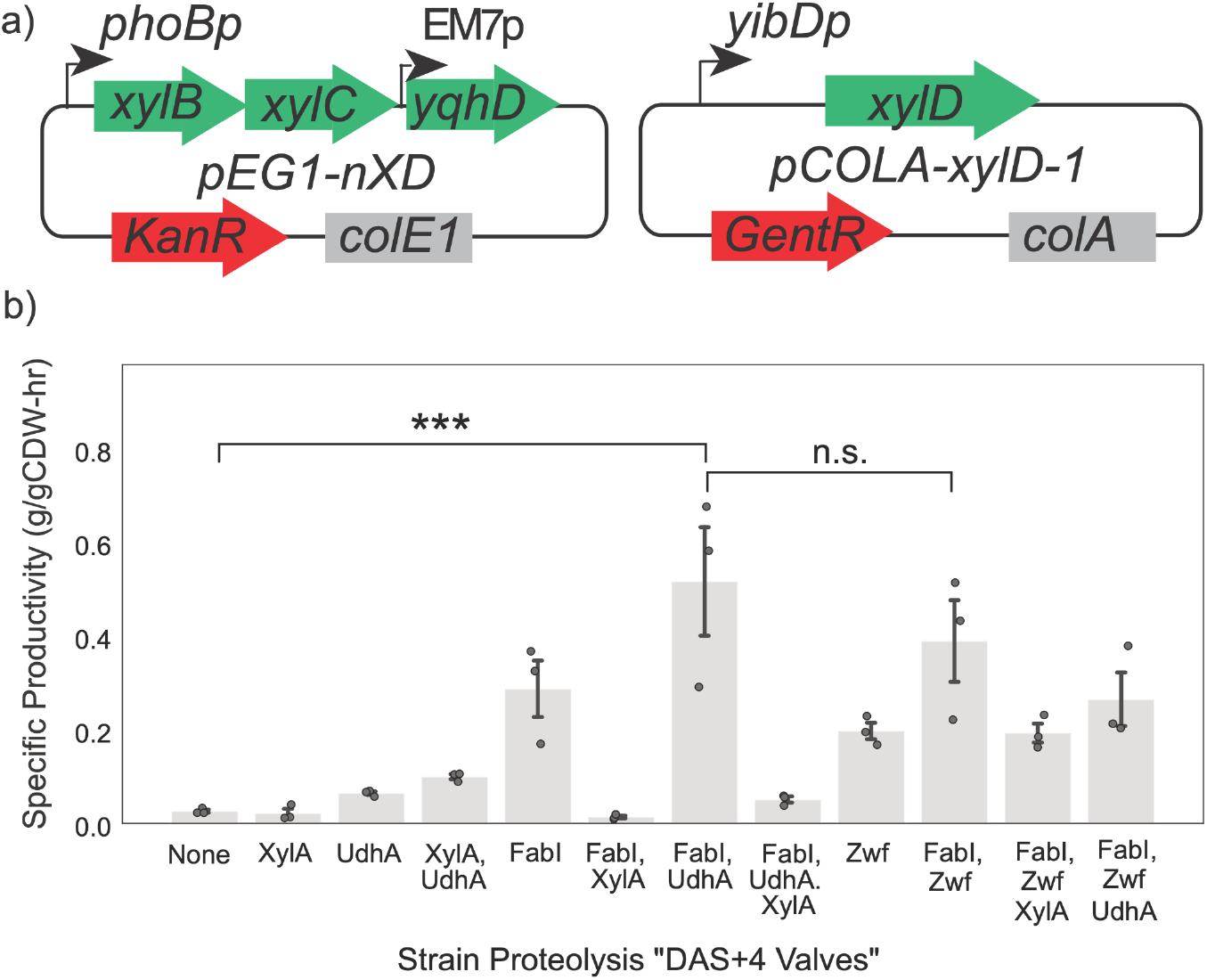
a) Maps of plasmids pEG1-nXD and pCOLA-xylD-1. b) Specific EG production (g/gCDW-hr) in microfermentations of strains with plasmids pEG1-nXD and pCOLA-xylD-1 in the control host DLF_Z0025 (No valves) as well as in strains with dynamic proteolysis of XylA, UdhA, FabI and Zwf alone and or in combination. A triple asterisk indicates a p-value <0.001, using a Welch’s t-test, comparing the strain with dynamic proteolysis of both FabI and UdhA to the control. No significant difference (n.s.) was measured between the strain with FabI and UdhA valves and the next highest producer with valves in FabI and Zwf.

We chose to move forward with our first variant, pCOLA-xylD-1, having the highest expression levels via SDS-PAGE. We transformed our panel of strains with plasmids pEG-1 and pCOLA-xylD-1 and evaluated these strains for EG production. Results are given in Figure 4b. Interestingly, with the improved XylD expression levels, the regulatory strains containing FabI and Zwf valves had a significant improvement in specific EG production rates achieving >0.5 g/gCDW-hr. It is worth noting that these results rely on dynamic control implemented only using proteolysis based degron tags. We have previously reported that the addition of CRISPR based gene silencing improved reductions in several enzymes including Zwf and FabI when compared to proteolysis alone. ^21,28,30^ We did evaluate strain variants with gene silencing of *zwf*, however none out performed the use of degron tags alone (Supplemental Figure S4). *FabI* silencing guides were not tested in this strain lineage due to reported stability issues.^42^

Finally, based on our microfermentation results, we evaluated several strains in instrumented bioreactors under minimal media fedbatch conditions. ^60^ The target biomass levels produced in the growth stage were ∼25 gCDW/L. Results are given in Figure 5. As cells approached their last doubling from OD_600nm_∼30 to OD_600nm_∼ 60, cells entered the stationary production stage where the EG biosynthesis pathway enzymes are induced and valve enzymes are “turned off”. The control strain with no valves bearing the expression plasmid pEG-1 produced ∼20g/Lof EG (Figure 5a). As seen in Figure 5b, strain PS_0001, with dynamic control over UdhA and XylA activity, also expressing plasmid pEG-1, produced ∼60 g/L of EG in 70 hours of production, with a yield of 0.3g ethylene glycol /g xylose (which is 72.6% of the theoretical). Lastly, as shown in Figure 5c, strain DLF_Z0040, with dynamic control over both FabI and Zwf activity, bearing plasmids pEG1-nXD and pCOLA-xylD-1, produced 140g/L EG, in 70 hours, with a yield of ∼0.38g EG /g xylose or 92% of the theoretical yield.

**Figure 5.**
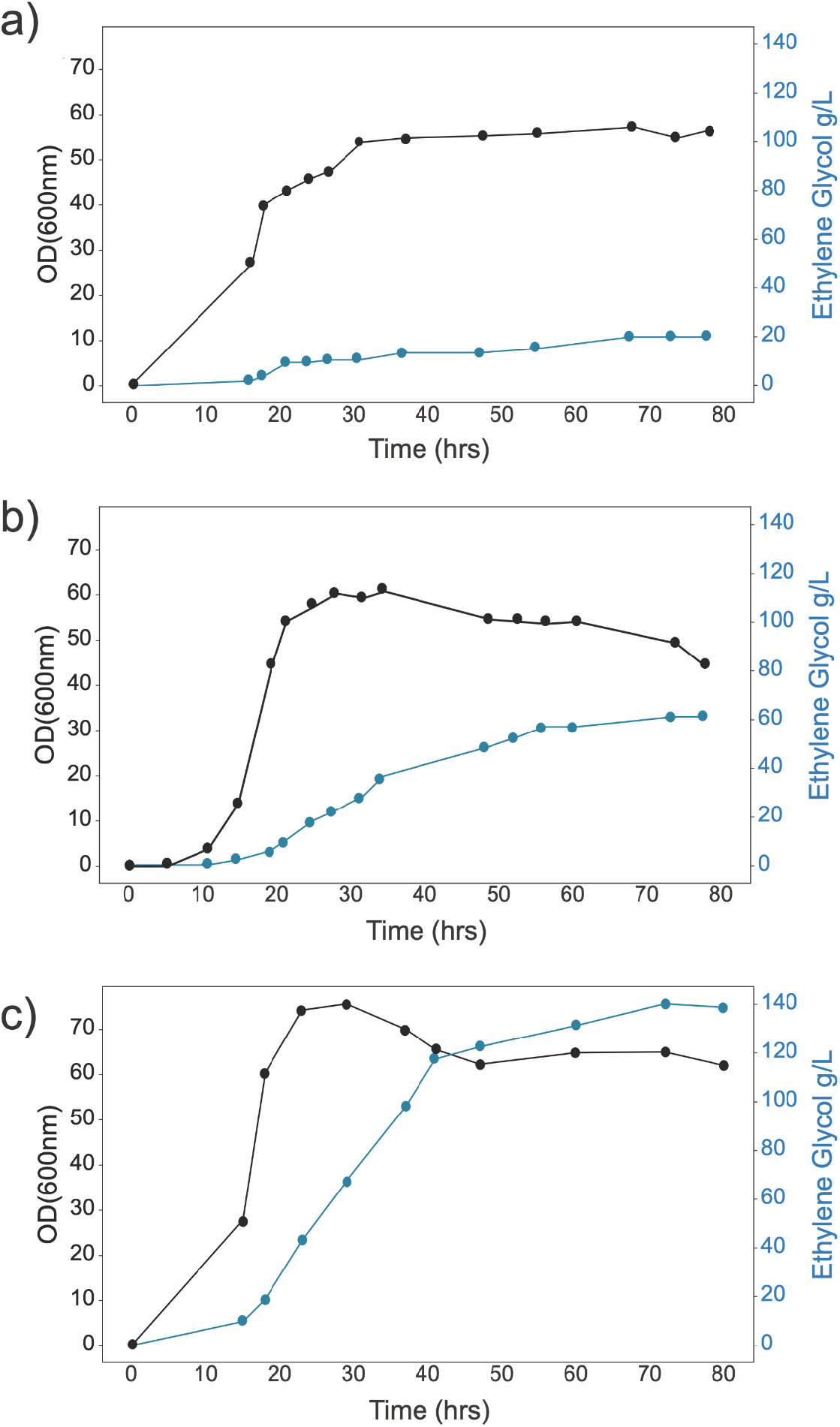
a) EG production in fed-batch fermentations in instrumented bioreactors. Biomass levels (black, OD600nm) and ethylene glycol levels (g/L, blue) are given as a function of time. a) Production via the control strain (DLF_Z0025, pEG-1), b) Production via the “UdhA, XylA” proteolysis valve strain PS_0001 plus pEG-1. c) Production via the “FabI, Zwf” proteolysis valve strain PS_0006 plus pEG1-nXD plus pCOLA-xylD-1 in fed batch fermentation shown in panel C, xylose feed rates were increased from 4g/hr to 8g/hr at 36 hrs, due to depletion of xylose.

## Discussion

Our results highlight the importance of identifying and overcoming multiple limiting fluxes in engineered metabolic networks to achieve high-level production, in this case of EG. Our data is consistent with two key limiting fluxes in the initial strains leveraging plasmid pEG-1: the carbon flux towards a competitive pathway (xylulose), the NADPH flux to a competitive pathway (NADH). Our initial stoichiometric valves, which dynamically degrade the competitive enzymes XylA (xylose isomerase) and UdhA (soluble transhydrogenase) in combination, resulted in a >3-fold improvement in specific productivity, reaching 0.11 g/gCDW-hr in microfermentations. This demonstrated a synergistic effect of rerouting both carbon and NADPH to the desired product. However, additional valves aimed at improving NADPH flux had no positive impact on EG production with our initial pathway expression construct. This is consistent with the hypothesis that these best “stoichiometric strains” were now limited by pathway enzyme activity rather than substrate availability. When we re-engineered the pathway by expressing XylD on a separate, optimized plasmid, pCOLA-xylD-1. Our two-plasmid system (pEG1-nXD and pCOLA-xylD-1) ensured robust expression of all four essential pathway enzymes, but expression alone did not improve EG biosynthesis. However, overcoming limitations in pathway activity as well as supplying additional NADPH flux using a regulatory strategy led to synergistic improvements in EG production, which supports the idea that improved pathway variants were now limited by NADPH flux rather than competitive pathways. ethylene glycol biosynthesis, as well as regulatory pathways aimed at increasing NADPH fluxes. Together, these data support a central hypothesis that faster pathways by their inherent speed can compete with other metabolic fluxes. In short the faster a pathway flux, the less impact a competitive flux will have. Additionally, faster pathways are better aided by regulatory changes capable of improving rates of substrate or cofactor synthesis, rather than removing competitive consumption. These results indicate that network modifications that are optimal for a slower pathway are not necessarily optimal for a faster pathway, and thus optimization strategies should not “lock in” modifications that may become suboptimal as the pathway’s speed is increased. This study thus provides a compelling demonstration of how pathway speed dictates the most effective strategy offering a new framework for metabolic engineering.

Lastly, our findings from microfermentations were successfully translated to instrumented fed-batch bioreactors, which are essential for evaluating industrial potential. The most significant achievement was with the strain containing the FabI and Zwf regulatory valves, combined with the optimized two-plasmid system, which produced an impressive 140 g/L of EG in 70 hours, achieving a yield of ∼0.38 g EG/g xylose, or 92% of the theoretical maximum. To our knowledge this result is the highest reported EG titer demonstrated in *E. coli* and highlights the power of dynamic control and systematic pathway optimization to achieve high titers and yields. Our study provides a clear roadmap for identifying and overcoming multiple limiting metabolic fluxes in a stepwise manner to significantly enhance bioproduction.

## Methods

### Reagents and Media

All reagents and materials were purchased from Sigma Aldrich except as noted below. MOPS (3-(N-morpholino) propanesulfonic acid) was obtained from BioBasic, Inc. (Amherst, NY). Crystalline xylose was obtained from Profood International (Naperville, IL). SM10++, SM10 No Phosphate, FGM10 and FGM25 media were prepared as previously reported. ^20,22,43,44^ However, glucose in these formulas was replaced by xylose at the same proportion (1g xylose for 1 g glucose) Low salt Luria Broth Lennox formulation was used for routine strain culture and construction. Working antibiotic concentrations were as follows: chloramphenicol: 35 µg/mL, kanamycin: 35 µg/mL, blasticidin: 100 µg/mL, ampicillin 50 µg/mL, gentamicin:50 µg/mL, zeocin: 100 µg/mL.

### Strains and Plasmids

All bacterial strains used in this study were generated using standard genetic engineering methodologies as previously described.^20,43^ A complete list of strains and plasmids used in this study is provided in Table 1, and the sequences of synthetic DNA fragments are listed in Supplemental Table S2. Chromosomal modifications were generated using standard recombineering methodologies.^45^ The recombineering plasmid pSIM5 was generously provided by Donald Court (NCI; https://redrecombineering.ncifcrf.gov/court-lab.html).^45,46^ C-terminal DAS+4 tags were introduced by direct chromosomal integration, with selection mediated through the integration of antibiotic resistance cassettes downstream of the target genes, *xylA, udhA, fabI*, and *zwf* in strain DLF_Z0025 as previously described.^20^ All constructed strains were validated by PCR, agarose gel electrophoresis, and sequencing of PCR amplicons. The complete list of oligonucleotides used for strain confirmation and sequencing is provided in Supplemental Table S3.

The plasmid pEG-1 used in this study was assembled from a synthetic linear DNA fragment (Gblocks^™^; Integrated DNA Technologies, Coralville, IA). This construct employed low-phosphate-inducible promoters, and the complete sequence is provided in Supplemental Table S2. To generate pEG-1, the Gblock containing xylB, xylC, xylD, and yqhD gene sequences was cloned into a linearized pSMART vector backbone via Gibson assembly. To construct pEG1-nXD, DNA fragments containing *xylB* and *xylC*, or *yqhD* were PCR-amplified. These two fragments, containing the promoter-gene cassettes, were integrated into a linearized pSMART backbone with overlapping ends using Gibson assembly. To construct pCOLA-xylD-1, the *xylD* gene sequence with an upstream yibDp promoter was PCR-amplified from pEG-1 and inserted into a linearized pCOLA vector backbone to construct pCOLA-xylD-1. To introduce His-tag and HiBiT-tag sequences at the N- and C-termini of xylD variants, primers incorporating the respective tags with overlapping ends were employed in Gibson assembly to construct pCOLA-xylD-3, pCOLA-xylD-4, and pCOLA-xylD-5. All plasmid constructs were verified by PCR, agarose gel electrophoresis, and confirmed through complete plasmid sequencing.

### Microfermentations

Microfermentation protocols were performed as previously described.^20–22,29,43,44,47^ Briefly, strains were propagated in LB Lennox (Low salt) overnight. On the second day, 5 μL of each strain was inoculated into 145μL SM10++ media with xylose as substrate and appropriate antibiotics in flat bottom 96 well plates in triplicate. Next, these cultures were grown for 18-24 hours at 37°C in a 300 rpm shaker, using the System Duetz or until mid exponential phase was obtained, defined as an OD600nm in the range of 5-15.^29^ Cells were then harvested by centrifugation at 3500 rpm for 10 mins, then pellets were washed with SM10 no-phosphate media by resuspending and centrifuging at 3500 rpm for 10 mins, to remove phosphate. Next, the washed pellets were resuspended in SM10 no-phosphate and simultaneously normalized biomass levels (OD_600nm_ = 1). Subsequently, cultures were again incubated in 96-well flat bottom plates at 37 °C and 300 rpm for 24 hours. After 24 hours, cells were harvested by centrifugation and supernatant collected for analytical assays.

### Seed Preparation

A 5 mL LB culture, supplemented with the appropriate antibiotics, was inoculated with the specific strain from a glycerol stock and incubated at 37°C overnight. Subsequently, 1% of the overnight culture was transferred into a 250 mL baffled shake flask containing 50 mL of SM10++ media with xylose as substrate and appropriate antibiotics and incubated at 37°C overnight. When the culture’s OD600 nm reached 6-10, a 50 mL culture of SM10 media (containing half the concentration of casamino acids and yeast extract as SM10++) was inoculated with 1% of the SM10++ culture and incubated under the same conditions. Again, when the culture’s OD600 nm reached 6-10, a 50 mL culture of SM10 media (without casamino acids or yeast extract) was inoculated similarly. Cells were harvested when the culture’s final OD600 nm reached 6-10. The culture was centrifuged at 4,000 rpm for 15 minutes to remove the media, and the harvested cells were resuspended in FGM10 media to achieve an OD600 nm of 10. Seed vials were created by mixing 6.5 mL of the resuspended culture with 1.5 mL of sterile 50% Glycerol for long-term storage at -70°C.

### 2-Stage Fermentations in Instrumented Bioreactors

A parallel bioreactor system (Infors-HT Multifors, Laurel, MD, USA) was employed for conducting the fermentation experiments. Briefly, tanks were filled with 800 mL FGM25 media formulation as previously reported ^43^ containing an appropriate phosphate concentration to achieve a final E. coli biomass of approximately 25 gCDW/L. Antibiotics were added as needed. Essential components, including phosphate, acidified ferrous sulfate, xylose, thiamine, and antibiotics, were introduced after sterilizing and cooling the tank vessel containing the remaining media components. Frozen seed vials containing 8 mL of seed culture were used to inoculate the tanks. Temperature and pH within the bioreactors were closely regulated at 37 °C and 6.8, respectively, using 14.5 M ammonium hydroxide and 1 M hydrochloric acid as titrants. The reactor was equipped with a PID control system, which maintained control over the dissolved oxygen concentration of 25%. Concentrated sterile filtered xylose feed (500 g/L) was added to the tanks at an initial rate of 3.5 g/L-h when the rate of agitation increased above 1000 rpm. This rate was then increased exponentially, until 40 g total glucose had been added, at which point the feed was set to 1g/L-hr. The starting batch xylose concentration was 25 g/L for both the tanks. Alternatively, for 1 L fed-batch fermentation of the best ethylene glycol producing strain in FGM25, tanks were initially filled with 650 mL medium containing 60 g L^−1^ xylose. Temperature and pH were maintained at 37 °C and pH 6.8 using 10 M NH_4_OH and 1 M HCl. A sterile-filtered xylose feed (600 g L^−1^) was fed under PID control with an exponential rate until 100 mL (80 g) feed was added when agitation exceeded 1000 rpm. The feed was then maintained at 4 g h^−1^ and increased to 8 g h^−1^ after 37 h to maximize ethylene glycol production.

### Expression Analysis & Densitometry

Expression of pEG-1, pEG1-nXD, pCOLA-xylD-1, pCOLA-xylD-3, pCOLA-xylD-4, and pCOLA-xylD-5 was carried out in the autolysis strain DLF_R004 using the microtiter plate expression and autolysis protocol previously described.^39^ Protein expression levels were quantified by densitometric analysis of SDS–PAGE gels. Whole-cell samples were normalized to OD600=10, mixed 1:1 with 2× loading buffer, and heated at 95 °C for 10 min. A 10 µL aliquot of each sample was loaded onto a 4–15% gradient Mini-PROTEAN TGX precast gel (Bio-Rad Laboratories, Hercules, CA) and ran at 160 V. Densitometry was performed using FIJI.^48,49^

### Analytical Methods

High performance liquid chromatography with a refractive index detector was used to RI quantify both ethylene glycol and xylose as described in previous work.^24^ Briefly, to separate xylose, ethylene glycol and other intermediates, we used a Rezex ROA-Organic Acid H^+^ (8%) HPLC Column (Cat #: #00H-0138-K0, Phenomenex, Inc., Torrance, CA, 300 x 7.8 mm) and a H-Class UPLC (Waters Acquity) with a 2414 Refractive Index (RI) detector (Waters Corp., Milford, MA. USA). MassLynx v4.1 software was employed for all analyses. An isocratic mobile phase consisting of 5 mM sulfuric acid was utilized at a flow rate of 0.5 mL/min, with a column temperature set at 50 °C. The injection volume of samples and standards was 10 μL. In order to maintain linearity within the analytical range, the samples were diluted 5-fold for microfermentation samples and 20-80 folds for bioreactor samples, in water.

## Author contributions

P. Sarkar and S. Li constructed plasmids and strains, performed microfermentations and instrumented fermentations and analytical analyses. J Chen performed fermentation experiments. U. Yano performed microfermentations, expression analysis and analytical analyses. M.D. Lynch designed experiments. All authors analyzed results, wrote, revised and edited the manuscript.

## Acknowledgements

We would like to acknowledge the following support: ONR YIP #12043956, SERDP WP20-1391, and NAVAIR contract # N6893623P0295.

