## Supplemental Materials for "Improved Biosynthesis of Ethylene Glycol from Xylose in Engineered *E. coli* Utilizing Two-Stage Dynamic Control"

**Table S1:** Reported kinetics for key enzymes involved in EG Biosynthesis

| Enzyme | Source (Gene) | Specific Activity (µmol/min-mg) | Kcat (s-1) | Km (µM) | Ref. |
|---|---|---|---|---|---|
| Xylose dehydrogenase | *C. crescentus xylB* | 58.1** | 25.8 | 400 | 1 |
| Xylonolactonase | *C. crescentus xylC* | >50* (estimate) | – | – | 2 |
| Xylonate dehydratase | *E. coli yjhG* | 0.014 | – | – | 3 |
| | *E. coli yagF* | 0.034 | – | – | 3 |
| | *C. crescentus xylD* | 21.8 | 23.43 | 1900 | 4 |
| Aldehyde reductase | *E. coli yqhD* | 76.9** | 54 | 28280 | 5 |
| *estimate from literature, not measured<br>** estimated from kcat and protein MW | | | | | |

**Table S2:** Synthetic DNA utilized for strain construction.

| **xylA-DAS4-ampR** |
|---|
| GATACGATGGCACTGGCGCTGAAAATTGCAGCGCGCATGATTGAAGATGGCGAGCTGGATAAACGCATCGCGC<br>AGCGTTATTCCGGCTGGAATAGCGAATTGGGCCAGCAAATCCTGAAAGGCCAAATGTCACTGGCAGATTTAGC<br>CAAATATGCTCAGGAACATCATTTGTCTCCGGTGCATCAGAGTGGTCGCCAGGAACAACTGGAAAATCTGGTAA<br>ACCATTATCTGTTCGACAAAGCGGCCAACGATGAAAACTATTCTGAAAACTATGCGGATGCGTCTTAATGATAAG<br>GACCGTGTTGACAATTAATCATCGGCATAGTATATCGGCATAGTATAATACGACAAGGTGAGGAACTAAACCATG<br>AGTATTCAACATTTCCGTGTCGCCCTTATTCCCTTTTTTGCGGCATTTTGCCTTCCTGTTTTTGCTCACCCAGAA<br>ACGCTGGTGAAAGTAAAAGATGCTGAAGATCAGTTGGGTGCACGAGTGGGTTACATCGAACTGGATCTCAACA<br>GCGGTAAGATCCTTGAGAGTTTACGCCCCGAAGAACGTTTTCCAATGATGAGCACTTTTAAAGTTCTGCTATGT<br>GGCGCGGTATTATCCCGTATTGACGCCGGGCAAGAGCAACTCGGTCGCCGCATACACTATTCTCAGAATGACT<br>TGGTTGAGTACTCACCAGTCACAGAAAAGCATCTCACGGATGGCATGACAGTAAGAGAATTATGCAGTGCTGC<br>CATAACCATGAGTGATAACACTGCGGCCAACTTACTTCTGGCAACGATCGGAGGACCGAAGGAGCTAACCGCT<br>TTTTTGCACAACATGGGGGATCATGTAACTCGCCTTGATCGTTGGGAACCGGAGCTGAATGAAGCCATACCAAA<br>CGACGAGCGTGACACCACGATGCCTGTAGCAATGGCAACAACGTTGCGCAAACTATTAACTGGCGAACTACTT<br>ACTCTAGCTTCCCGGCAACAATTAATAGACTGGATGGAGGCGGATAAAGTTGCAGGATCACTTCTGCGCTCGG<br>CCCTCCCGGCTGGCTGGTTTATTGCTGATAAATCTGGAGCCGGTGAGCGTGGGTCTCGCGGTATCATTGCAGC<br>ACTGGGGCCAGATGGTAAGCCCTCCCGCATCGTAGTTATCTACACGACGGGGAGTCAGGCAACTATGGATGAA<br>CGAAATAGACAGATCGCTGAGATAGGTGCCTCACTGATTAAGCATTGGTAGTAAGTAGGGATAACAGGGTAATC |

GGCTAACTGTGCAGTCCGTTGGCCCGGTTATCGGTAGCGATACCGGGCATTTTTTTAAGGAACGATCGATATGT
ATATCGGGATAGATCTTGGCACCTCGGGCGTAAAAGTTATTTTGCTCAACGAGCAGGGTGAGGTGGTTGCTGC
GCAAACGGAAAAGCTGACCGTTTCGCGCCCGCATCCACTCTGGTCGGAACAAGACCCGGAACAGTGGTGGC
AGGCAACTGATCGCGCAA

**udhA-DAS+4-bsdR**

TCTGGGTATTCACTGCTTTGGCGAGCGCGCTGCCGAAATTATTCATATCGGTCAGGCGATTATGGAACAGAAAG
GTGGCGGCAACACTATTGAGTACTTCGTCAACACCACCTTTAACTACCCGACGATGGCGGAAGCCTATCGGGT
AGCTGCGTTAAACGGTTTAAACCGCCTGTTTGCGGCCAACGATGAAAACTATTCTGAAAACTATGCGGATGCGT
CTTAATAGTTGACAATTAATCATCGGCATAGTATATCGGCATAGTATAATACGACTCACTATAGGAGGGCCATCATG
AAGACCTTCAACATCTCTCAGCAGGATCTGGAGCTGGTGGAGGTCGCCACTGAGAAGATCACCATGCTCTATG
AGGACAACAAGCACCATGTCGGGGCGGCCATCAGGACCAAGACTGGGGAGATCATCTCTGCTGTCCACATTG
AGGCCTACATTGGCAGGGTCACTGTCTGTGCTGAAGCCATTGCCATTGGGTCTGCTGTGAGCAACGGGCAGA
AGGACTTTGACACCATTGTGGCTGTCAGGCACCCCTACTCTGATGAGGTGGACAGATCCATCAGGGTGGTCAG
CCCCTGTGGCATGTGCAGAGAGCTCATCTCTGACTATGCTCCTGACTGCTTTGTGCTCATTGAGATGAATGGCA
AGCTGGTCAAAACCACCATTGAGGAACTCATCCCCCTCAAGTACACCAGGAACTAAAGTAAAACTTTATCGAAA
TGGCCATCCATTCTTGCGCGGATGGCCTCTGCCAGCTGCTCATAGCGGCTGCGCAGCGGTGAGCCAGGACGA
TAAACCAGGCCAATAGTGCGGCGTGGTTCCGGCTTAATGCACGG

**fabI-DAS+4-zeoR**

atgGGTTTTCTTTCCGGTAAGCGCATTCTGGTAACCGGTGTTGCCAGCAAACTATCCATCGCCTACGGTATCGCT
CAGGCGATGCACCGCGAAGGAGCTGAACTGGCATTCACCTACCAGAACGACAAACTGAAAGGCCGCGTAGAA
GAATTTGCCGCTCAATTGGGTTCTGACATCGTTCTGCAGTGCGATGTTGCAGAAGATGCCAGCATCGACACCAT
GTTCGCTGAACTGGGGAAAGTTTGGCCGAAATTTGACGGTTTCGTACACTCTATTGGTTTTGCACCTGGCGATC
AGCTGGATGGTGACTATGTTAACGCCGTTACCCGTGAAGGCTTCAAAATTGCCCACGACATCAGCTCCTACAG
CTTCGTTGCAATGGCAAAAGCTTGCCGCTCCATGCTGAATCCGGGTTCTGCCCTGCTGACCCTTTCCTACCTT
GGCGCTGAGCGCGCTATCCCGAACTACAACGTTATGGGTCTGGCAAAAGCGTCTCTGGAAGCGAACGTGCGC
TATATGGCGAACGCGATGGGTCCGGAAGGTGTGCGTGTTAACGCCATCTCTGCTGGTCCGATCCGTACTCTGG
CGGCCTCCGGTATCAAAGACTTCCGCAAAATGCTGGCTCATTGCGAAGCCGTTACCCCGATTCGCCGTACCGT
TACTATTGAAGATGTGGGTAACTCTGCGGCATTCCTGTGCTCCGATCTCTCTGCCGGTATCTCCGGTGAAGTGG
TCCACGTTGACGGCGGTTTCAGCATTGCTGCAATGAACGAACTCGAACTGAAAGCGGCCAACGATGAAAACTA
TTCTGAAAACTATGCGGATGCGTCTtaaTTGACAATTAATCATCGGCATAGTATATCGGCATAGTATAATACGACTC
ACTATAGGAGGGCCATCATGGCCAAGTTGACCAGTGCCGTTCCGGTGCTCACCGCGCGCGACGTCGCCGGA
GCGGTCGAGTTCTGGACCGACCGGCTCGGGTTCTCCCGGGACTTCGTGGAGGACGACTTCGCCGGTGTGGT
CCGGGACGACGTGACCCTGTTCATCAGCGCGGTCCAGGACCAGGTGGTGCCGGACAACACCCTGGCCTGGG
TGTGGGTGCGCGGCCTGGACGAGCTGTACGCCGAGTGGTCGGAGGTCGTGTCCACGAACTTCCGGGACGC
CTCCGGGCCGGCCATGACCGAGATCGGCGAGCAGCCGTGGGGGCGGGAGTTCGCCCTGCGCGACCCGGCC
GGCAACTGCGTGCACTTTGTGGCAGAGGAGCAGGACTGATCGTTCTGTTGGTAAAGATGGGCGGCGTTCTGC
CGCCCGTTATCTCTGTTATACCTTTCTGATATTTGTTATCGCCGATCCGTCTTTCTCCCCTTCCCGCCTTGCGTC
AGGATAACGATTTCCTTTACGACCAAGGAGCGCCCATGGAACAACGCCACATCACCGGCAAAAGCCACTGGTA
TCATGAAACGCAATCCAGTACTACGGAGTATGACGTTCTGCCTCTGGTCCCGGAAGCCGCAAAGGTCAGCGAT
CCCTTTCTACTCGACGTGATCCTTGAAAAAGAAACGCTGGCCCCCTTCCTTTCATGGCTGGACCCTGCGCGTG
TTCTTGCAGTGGATTTGTTCCCTGACCAGCTTACCGTGACCCGT

**zwf-DAS+4-bsdR**

GAAGTGGAAGAAGCCTGGAAATGGGTAGACTCCATTACTGAGGCGTGGGCGATGGACAATGATGCGCCGAAA
CCGTATCAGGCCGGAACCTGGGGACCCGTTGCCTCGGTGGCGATGATTACCCGTGATGGTCGTTCCTGGAAT
GAGTTTGAGGCGGCCAACGATGAAAACTATTCTGAAAACTATGCGGATGCGTCTTAATAGTTGACAATTAATCAT
CGGCATAGTATATCGGCATAGTATAATACGACTCACTATAGGAGGGCCATCATGAAGACCTTCAACATCTCTCAG
CAGGATCTGGAGCTGGTGGAGGTCGCCACTGAGAAGATCACCATGCTCTATGAGGACAACAAGCACCATGTC

GGGGCGGCCATCAGGACCAAGACTGGGGAGATCATCTCTGCTGTCCACATTGAGGCCTACATTGGCAGGGTC
ACTGTCTGTGCTGAAGCCATTGCCATTGGGTCTGCTGTGAGCAACGGGCAGAAGGACTTTGACACCATTGTGG
CTGTCAGGCACCCCTACTCTGATGAGGTGGACAGATCCATCAGGGTGGTCAGCCCCTGTGGCATGTGCAGAG
AGCTCATCTCTGACTATGCTCCTGACTGCTTTGTGCTCATTGAGATGAATGGCAAGCTGGTCAAAACCACCATT
GAGGAACTCATCCCCCTCAAGTACACCAGGAACTAAAGTAATATCTGCGCTTATCCTTTATGGTTATTTTACCGG
TAACATGATCTTGCGCAGATTGTAGAACAATTTTTACACTTTCAGGCCTCGTGCGGATTCACCCACGAGGCTTTT
TTTATTACACTGACTGAAACGTTTTTGCCCTATGAGCTCCGGTTACAGGCGTTTCAGTCATAAATCCTCTGAATG
AAACGCGTTGTGAATC

**pEG-1 insert**

CATGCATAATCCGCACGCATCTGGAATAAGGAAGTGCCATTCCGCCTGACCTGCCACGGAAATCAATAACCTGA
AGATATGTGCGACGAGCTTTTCATAAATCTGTCATAAATCTGACGCATAATGACGTCGCATTAATGATCGCAACCT
ATTTATTGTGTAGGAGGATAATCTATGGCTAGCAAAGGAGAAGAACTTTTCACATGAGTAGCGCCATTTACCCAT
CTCTGAAGGGAAAGCGTGTCGTGATTACCGGGGGTGGCTCGGGAATTGGGGCTGGTTTAACAGCTGGTTTCG
CTCGTCAAGGTGCGGAAGTAATCTTCTTGGATATTGCTGATGAGGACTCCCGTGCTCTTGAAGCGGAACTTGC
CGGTAGCCCTATCCCACCCGTATACAAGCGTTGTGATCTGATGAACCTGGAAGCTATCAAAGCCGTTTTCGCTG
AGATTGGTGACGTTGATGTCTTGGTCAACAATGCGGGTAACGACGACCGCCATAAGTTGGCGGACGTCACTGG
AGCATATTGGGACGAGCGTATTAATGTCAATCTGCGCCATATGTTGTTTTGCACGCAGGCGGTGGCTCCGGGC
ATGAAAAAACGCGGGGGAGGGGCAGTCATTAATTTTGGAAGCATCAGCTGGCACCTTGGCTTGGAAGATCTTG
TTTTATATGAAACTGCCAAGGCTGGGATTGAGGGGATGACTCGTGCTCTGGCACGTGAGTTGGGTCCAGACGA
CATTCGCGTCACATGCGTTGTCCCCGGTAATGTCAAGACTAAACGCCAAGAGAAATGGTATACCCCTGAGGGG
GAAGCGCAGATTGTGGCCGCGCAATGCCTTAAAGGCCGCATCGTACCAGAAAATGTTGCAGCGCTGGTTTTAT
TCTTAGCAAGCGATGACGCCAGTTTATGTACAGGGCATGAATACTGGATCGACGCCGGATGGCGCTGATAAGG
ATCTAGGAGGGAGATCATATGACCGCTCAAGTTACGTGTGTTTGGGACCTTAAGGCGACACTGGGTGAAGGTC
CGATCTGGCACGGCGATACCTTATGGTTTGTCGATATTAAGCAGCGTAAAATCCACAACTACCACCCTGCGACT
GGAGAGCGTTTTTCGTTCGACGCGCCCGATCAAGTGACTTTTTTAGCACCGATTGTTGGAGCGACTGGGTTTG
TCGTCGGCTTGAAGACTGGCATTCATCGCTTTCACCCAGCGACTGGTTTCTCGTTGTTGTTAGAAGTCGAGGA
TGCCGCATTGAATAACCGCCCCAATGATGCTACTGTCGATGCCCAGGGTCGCCTTTGGTTCGGGACCATGCAC
GATGGAGAGGAAAACAACAGCGGGTCGCTGTATCGCATGGACCTGACAGGAGTAGCCCGCATGGACCGTGAC
ATTTGCATTACTAATGGTCCATGCGTTTCCCCAGACGGTAAGACGTTTTACCACACGGATACACTTGAGAAGACT
ATTTATGCTTTCGACTTAGCCGAGGATGGACTGCTTTCAAACAAGCGTGTGTTCGTCCAATTTGCATTGGGAGA
TGATGTCTATCCCGATGGAAGTGTCGTAGATTCGGAGGGATACTTATGGACAGCCTTATGGGGCGGTTTTGGAG
CCGTCCGCTTTAGTCCTCAAGGTGATGCCGTTACACGCATCGAGCTGCCTGCCCCTAACGTCACAAAACCGTG
CTTCGGTGGCCCTGATCTGAAAACCCTTTATTTTACGACTGCTCGTAAGGGGTTATCGGATGAAACCCTTGCCC
AGTACCCACTGGCGGGGGGAGTGTTCGCAGTGCCGGTTGATGTGGCGGGTCAACCTCAACATGAGGTACGC
CTTGTCTAATAAGTGCGTAATTGTGCTGATCTCTTATATAGCTGCTCTCATTATCTCTCTACCCTGAAGTGACTCT
CTCACCTGTAAAAATAATATCTCACAGGCTTAATAGTTTCTTAATACAAAGCCTGTAAAACGTCAGGATAACTTCT
GTGTAGGAGGATAATCTATGTCGAACCGTACCCCGCGTCGTTTCCGTAGTCGTGATTGGTTTGACAATCCGGAT
CACATCGACATGACAGCGCTGTACTTAGAGCGTTTCATGAACTATGGCATTACTCCGGAAGAATTACGCAGTGG
CAAGCCCATTATCGGGATTGCGCAAACCGGCAGTGATATTAGTCCATGCAATCGCATTCACCTTGACTTAGTTCA
GCGTGTGCGTGACGGCATTCGTGATGCAGGCGGTATCCCAATGGAATTCCCGGTACATCCTATCTTCGAAAACT
GCCGCCGTCCCACAGCGGCGTTAGACCGTAATCTGAGTTACCTGGGGCTTGTTGAGACGTTACATGGGTACCC
TATTGACGCTGTTGTATTAACCACCGGTTGCGACAAAACAACGCCAGCTGGGATCATGGCAGCTACCACCGTC
AACATTCCAGCGATCGTATTATCAGGGGGTCCAATGTTAGATGGCTGGCACGAAAACGAGTTAGTTGGCTCCG
GCACGGTCATTTGGCGCTCACGTCGTAAACTGGCAGCTGGGGAAATTACAGAAGAAGAATTCATTGACCGTGC
TGCTTCTTCCGCTCCAAGCGCAGGCCACTGCAACACGATGGGAACTGCGTCTACGATGAATGCAGTGGCTGA
GGCTTTGGGCCTGTCTCTGACTGGGTGTGCAGCTATCCCAGCGCCTTATCGTGAACGCGGGCAGATGGCTTA
CAAGACGGGGCAGCGCATCGTTGACCTTGCGTATGACGACGTCAAGCCATTAGACATTTTAACTAAGCAGGCA
TTTGAAAACGCGATCGCGTTGGTCGCAGCCGCGGGTGGCAGTACGAATGCGCAGCCCCATATCGTTGCGATG
GCTCGTCATGCAGGGGTCGAAATCACAGCCGACGATTGGCGCGCTGCATACGACATTCCATTAATTGTTAATAT
GCAGCCTGCCGGCAAATACCTTGGGGAACGCTTTCACCGCGCTGGCGGCGCCCCTGCTGTCCTTTGGGAATT
ATTACAACAGGGGGCGTTTGCACGGTGACGTTTTAACCGTAACAGGGAAGACCATGTCAGAGAATTTGCAGGGA
CGCGAAACCTCTGACCGCGAAGTTATTTTTCCCTATCATGAGCCTCTTGCTGAGAAAGCAGGTTTCTTAGTTTT

| |
|---|
| GAAAGGAAACCTTTTCGACTTCGCCATCATGAAATCATCCGTAATTGGCGAAGAATTCCGTAAGCGCTATCTGA<br>GTCAACCCGGTCAAGAAGGTGTGTTCGAAGCCCGCGCTATTGTTTTTGATGGGTCAGATGATTATCACAAGCGT<br>ATTAACGACCCAGCACTGGAGATTGATGAACGTTGTATTTTAGTCATTCGTGGAGCTGGTCCGATTGGATGGCC<br>AGGATCTGCTGAAGTAGTGAATATGCAGCCTCCAGACCACCTGCTTAAGAAAGGTATTATGTCGCTGCCTACAC<br>TTGGTGACGGGCGTCAAAGCGGGACAGCGGACTCACCATCGATTCTGAACGCTTCTCCGGAGAGTGCCATCG<br>GCGGAGGCCTTTCGTGGCTGCGTACGGGCGATACGATTCGTATTGACTTAAATACAGGTCGCTGTGATGCTCT<br>GGTTGATGAAGCGACGATTGCGGCTCGCAAGCAAGACGGCATCCCAGCTGTGCCGGCAACAATGACTCCGTG<br>GCAGGAGATTTACCGTGCCCATGCATCGCAACTTGACACAGGTGGCGTACTGGAATTTGCGGTCAAGTATCAA<br>GATTTGGCGGCCAAACTTCCGCGTCATAATCATTAAGGACCGTGTTGACAATTAATCATCGGCATAGTATATCGG<br>CATAGTATAATACGACAAGGTGAGGAACTAAACCATGAACAACTTTAATCTGCACACCCCAACCCGCATTCTGTT<br>TGGTAAAGGCGCAATCGCTGGTTTACGCGAACAAATTCCTCACGATGCTCGCGTATTGATTACCTACGGCGGC<br>GGCAGCGTGAAAAAAACCGGCGTTCTCGATCAAGTTCTGGATGCCCTGAAAGGCATGGACGTGCTGGAATTT<br>GGCGGTATTGAGCCAAACCCGGCTTATGAAACGCTGATGAACGCCGTGAAACTGGTTCGCGAACAGAAAGTG<br>ACTTTCCTGCTGGCGGTTGGCGGCGGTTCTGTACTGGACGGCACCAAATTTATCGCCGCAGCGGCTAACTATC<br>CGGAAAATATCGATCCGTGGCACATTCTGCAAACGGGCGGTAAAGAGATTAAAAGCGCCATCCCGATGGGCTG<br>TGTGCTGACGCTGCCAGCAACCGGTTCAGAATCCAACGCAGGCGCGGTGATCTCCCGTAAAACCACAGGCGA<br>CAAGCAGGCGTTCCATTCTGCCCATGTTCAGCCGGTATTTGCCGTGCTCGATCCGGTTTATACCTACACCCTGC<br>CGCCGCGTCAGGTGGCTAACGGCGTAGTGGACGCCTTTGTACACACCGTGGAACAGTATGTTACCAAACCGG<br>TTGATGCCAAAATTCAGGACCGTTTCGCAGAAGGCATTTTGCTGACGCTAATCGAAGATGGTCCGAAAGCCCT<br>GAAAGAGCCAGAAAACTACGATGTGCGCGCCAACGTCATGTGGGCGGCGACTCAGGCGCTGAACGGTTTGAT<br>TGGCGCTGGCGTACCGCAGGACTGGGCAACGCATATGCTGGGCCACGAACTGACTGCGATGCACGGTCTGG<br>ATCACGCGCAAACACTGGCTATCGTCCTGCCTGCACTGTGGAATGAAAAACGCGATACCAAGCGCGCTAAGCT<br>GCTGCAATATGCTGAACGCGTCTGGAACATCACTGAAGGTTCCGATGATGAGCGTATTGACGCCGCGATTGCC<br>GCAACCCGCAATTTCTTTGAGCAATTAGGCGTGCCGACCCACCTCTCCGACTACGGTCTGGACGGCAGCTCC<br>ATCCCGGCTTTGCTGAAAAAACTGGAAGAGCACGGCATGACCCAACTGGGCGAAAATCATGACATTACGTTGG<br>ATGTCAGCCGCCGTATATACGAAGCCGCCCGCTAACAGGCTAGGTGGAGGCTCAGTGATGATAAGTCTGCGAT<br>GGTGGATGCATGTGTCATG |

**Table S3:** Oligos used in this study for plasmid and strain confirmations

| Oligo | Sequence |
|---|---|
| xylA_conf_F | AGATGGCGAGCTGGATA |
| ampR_intR | AGTACTCAACCAAGTCATTCTG |
| udhA_conf_F | CAAAAGAGATTCTGGGTATTCACT |
| bsdR_intR | GAGCATGGTGATCTTCTCAGT |
| fabID4-zeoR_500up | GGCCGAAATTTGACGGTTTCGTA |
| fabID4-zeoR_200dn | GGGGCCAGCGTTTCTTTTTC |
| zwf_conf_F | CTGCTGGAAACCATGCG |
| pCOLA_FP | GTTTAAACCGACAAGGTGAGGAATCCgagtcacactg |
| pCOLA_xylD_HIBIT_seq_R | TCACCTTGTCGGTTTAAACTCAGCTAATCTTCTTAAACAG |
| pCOLA_RP | ACGCAATTAATGTAAGTTAGCTCACTCATTAGGCACCGGG |

| | |
|---|---|
| pSMART_conf_fwd | ACGCAATTAATGTAAGTTAGCTCACTCATTAGGCACCGGG |
| pSMART_conf_rev | CAGGAAACAGCTATGAC |
| pEG1-nXD_phoB_F | CTGGAATAAGGAAGTGCCATTCCGCCTGACCTGCCACGGAAATCAATAAC |
| pEG1-nXD_xylC_R | ATACTATGCCGATGATTAATTGTCAACACGGTCCCTTATTAGACAAGGCGTA |
| pEG1-nXD_EM7_F | AGGTACGCCTTGTCTAATAAGGGACCGTGTTGACAATTAATCATCGGCATA |
| pEG-nXD_yqhD_R | ACTTATCATCACTGAGCCTCCACCTAGCCTGTTAGCGGGCGGCTTCGTAT |

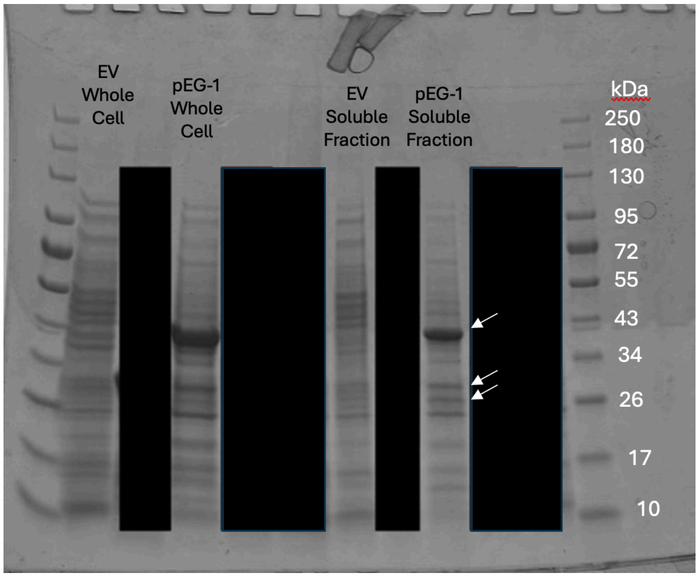


**Figure S1**. SDS-PAGE Expression analysis for plasmid pEG-1. EV: Empty vector control (pSMART-HC-Kan, Lucigen), plasmid not expressing any protein). Whole cell protein levels in strain R004 expressing plasmid pEG-1, after 24 hours of induction by phosphate depletion. Densitometry estimates for each protein are yqhD (42.1 kDa) 27.6%, xylC (31.6kDa) 8.75%, xylB (26.6kDa) 5.08%, overexpression of xylD (64.2 kDa) is not observed. Soluble expression levels in strain R004 with plasmid pEG-1, after 24 hours of induction by phosphate depletion. Densitometry estimates for each protein are yqhD (42.1 kD) 33.1%, xylC (31.6kD) 8.09%, xylB (26.6kD) 8.55%. Again expression of xylD (64.2 kDa) is not observed. Black boxes are used to obscure additional expression analysis not relevant to this study.

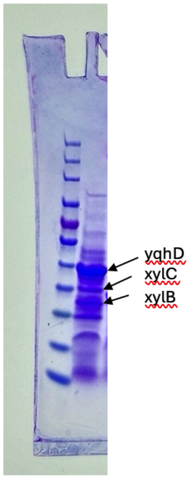


**Figure S2**. SDS-PAGE Expression analysis for plasmid pEG-1. EV: Empty vector control (pSMART-HC-Kan, Lucigen), plasmid not expressing any protein). Whole cell protein levels in strain R004 expressing plasmid pEG1-nXD, after 24 hours of induction by phosphate depletion. Densitometry estimates for each protein are yqhD (42.1 kDa) 19.6%, xylC (31.6kDa) 7.1%, xylB (26.6kDa) 10.3%.

**Table S4:** Overview of xylD variants

| **xylD variant** | **RBS** | **Coding sequence** |
|---|---|---|
| v1 | TGTGTAGGAGGATAATCT | MSNRTPR……HNH-Stop |
| v3 | ATATCCAAAACAGGAATAGCAGA AATAAGGAGGTATTTTT | MLNRTIAKRTLSNRTPR……HNH..linker-HIBIT-Stop |
| v4 | TGTGTAGGAGGATAATCT | MHHHHHHSNRTPR……HNH..linker-HIBIT-Stop |
| v5 | TGTCTGGATTCAGCAAGGAGGT TTTTTC | MHHHHHHSNRTPR……HNH..linker-HIBIT-Stop |
| …… " identical protein sequence" | | |

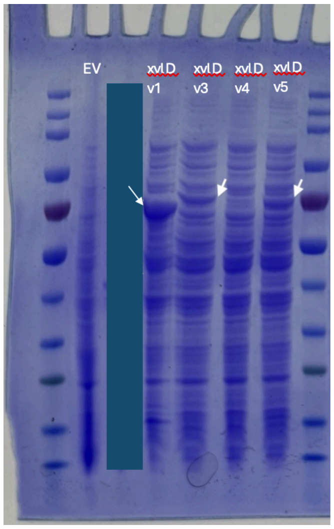


**Figure S3**. SDS-PAGE expression analysis for XylD expression variants: pCOLA-xylD-1, pCOLA-xylD-3, pCOLA-xylD-4, pCOLA-xylD-5. Whole cell protein levels in strain R004 expressing each plasmid after 24 hours of induction by phosphate depletion. Densitometry estimates for each protein are variant 1: 19.1%, variant 3: 6.82%, variant 4:  1.5% and variant 7.94 %.

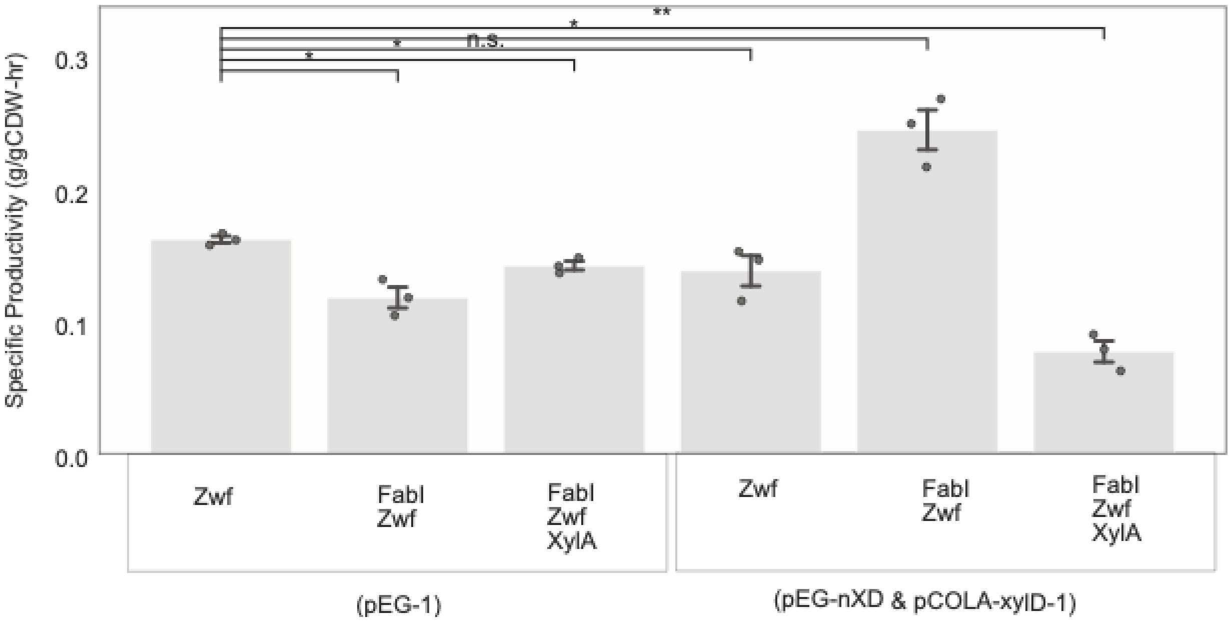


**Figure S4**. Specific EG production (g/gCDW-hr) in microfermentations of strains with plasmid pCASCADE-zwf, used for the dynamic silencing of *zwf* expression. Proteolysis valves in host strains include labels: Zwf, FabI and XylA. The left three bars are for pathway enzyme expression using plasmid pEG-1 and the right three bars are for pathway enzyme expression using plasmids pEG1-nXD and pCOLA-xylD-1.
